# Quantification of Focal Outflow Enhancement using Differential Canalograms

**DOI:** 10.1101/044503

**Authors:** Ralitsa T. Loewen, Eric N. Brown, Gordon Scott, Hardik Parikh, Joel S. Schuman, Nils A. Loewen

## Abstract

**Purpose:** To quantify regional changes of conventional outflow caused by ab interno trabeculectomy (AIT).

**Methods:** Gonioscopic, plasma-mediated ab interno trabeculectomy (AIT; Trabectome, Neomedix, Tustin, CA) was established in enucleated pig eyes. We developed a program to automatically quantify outflow changes (R, package eye-canalogram, github.com) using a fluorescent tracer reperfusion technique. Trabecular meshwork (TM) ablation was demonstrated with fluorescent spheres in 6 eyes before formal outflow quantification with two dye reperfusion canalograms in 6 further eyes. Eyes were perfused with a central, intracameral needle at 15 mmHg. Canalograms and histology were correlated for each eye.

**Results:** The pig eye provided a model with high similarity to AIT in human patients. Histology indicated ablation of TM and unroofing of most Schlemm’s canal segments. Spheres highlighted additional circumferential and radial outflow beyond the immediate area of ablation. Differential canalograms showed that AIT caused an increase of outflow of 17±5 fold inferonasally (IN), 14±3 fold superonasally (SN) and also an increase in the opposite quadrants with a 2±1 fold increase superotemporally (ST) and 3±3 inferotemporally (IT). Perilimbal specific flow image analysis showed an accelerated nasal filling with an additional perilimbal flow direction into adjacent quadrants.

**Conclusion:** A quantitative, differential canalography technique was developed that allows to quantify supraphysiological outflow enhancement by AIT.

## Introduction

Bulk outflow of aqueous humor and its relationship to intraocular pressure (IOP) has been well characterized.^1,2^ In contrast, focal outflow and enhancement of focal outflow by microincisional glaucoma surgeries has only been modeled mathematically^3,4^ but not been measured directly. Tracers have been used to highlight areas of increased flow through the TM where they become lodged near collector channel openings.^5,6^ The TM and the aqueous spaces of Schlemm’s canal and collector channels in the superficial to mid-level sclera can also be imaged non-invasively by spectral domain optical coherence tomography (SD-OCT).^7^ Wang et al used a Doppler strategy to detect movements of gold nanorods by SD-OCT but a quantification of flow could not be obtained.^8^ This approach may have limited use in vivo due to toxicity.^9^

It is estimated that the smallest of the current trabecular meshwork micro-bypass implants is limited to drainage segments of about 60 degrees,^10,11^ while larger ones access additional clock hours.^12^ Trabecular meshwork ablation can provide more extensive angle access by allowing to skip areas of discontinuity of Schlemm’s canal (SC).^11,13^ Although the trabecular meshwork is thought to be the anatomic location of primary outflow resistance, ^14^ intraocular pressures that are close to the theoretical limit of episcleral venous pressure^11^ are rarely achieved with trabecular bypasses^15^ or with trabecular ablation.^16^

Here, we hypothesized that it is possible to develop a differential canalography technique to directly analyze areas of altered outflow before and after an intervention. We refined recently introduced methods of quantitative canalography^17^ and used them to measure conventional outflow enhancement following plasma-mediated ab interno trabeculectomy.

## Methods

### Trabectome-Mediated Ab Interno Trabeculectomy in Pig Eyes

Pig eyes were obtained from a local abattoir. Only eyes that could be identified as right eyes were used. Within 2 hours of death eyelids and adnexal structures were excised, while the conjunctiva was preserved for the entire length of the globe. Eyes irrigated with phosphate buffered saline (PBS) were placed with the optic nerve into a cryogenic vial cup (CryoElite Cryogenic Vial #W985100, Wheaton Science Products, Millville, NJ) for compression free mount as described before.^17^ Similar to AIT in human eyes (Figure 1, A),^11^ eyes were positioned under a surgical microscope looking up with the temporal side of the eye directed towards the surgeon. A clear corneal incision was fashioned with a 1.8 mm keratome approximately 2 mm anterior to the temporal limbus while the inside was slightly flared for a striae free visualization during the procedure. Eyes were then tilted 30 degrees away from the surgeon and a goniolens (Trabectome Goniolens ONT-L, #600010, Neomedix Inc., Tustin, CA) was placed on the cornea to visualize the chamber angle (Figure 1, B). The tip of a Trabectome handpiece (handpiece #600018, Neomedix Inc., Tustin, CA) that was connected to a standard Trabectome system (Trabectome System #600026, Neomedix Inc., Tustin, CA) was inserted and advanced to the opposite chamber angle. After gentle goniosynechiolysis with the side of the instrument’s tip to disengage pectinate ligaments, TM ablation was continued towards the left by 45 degrees. The tip was then turned around inside of the eye, the TM was engaged again, and ablation continued by 45 degrees in the opposite direction. The instrument was withdrawn and the incision sealed with a drop of cyanoacrylate.

**Figure 1:**
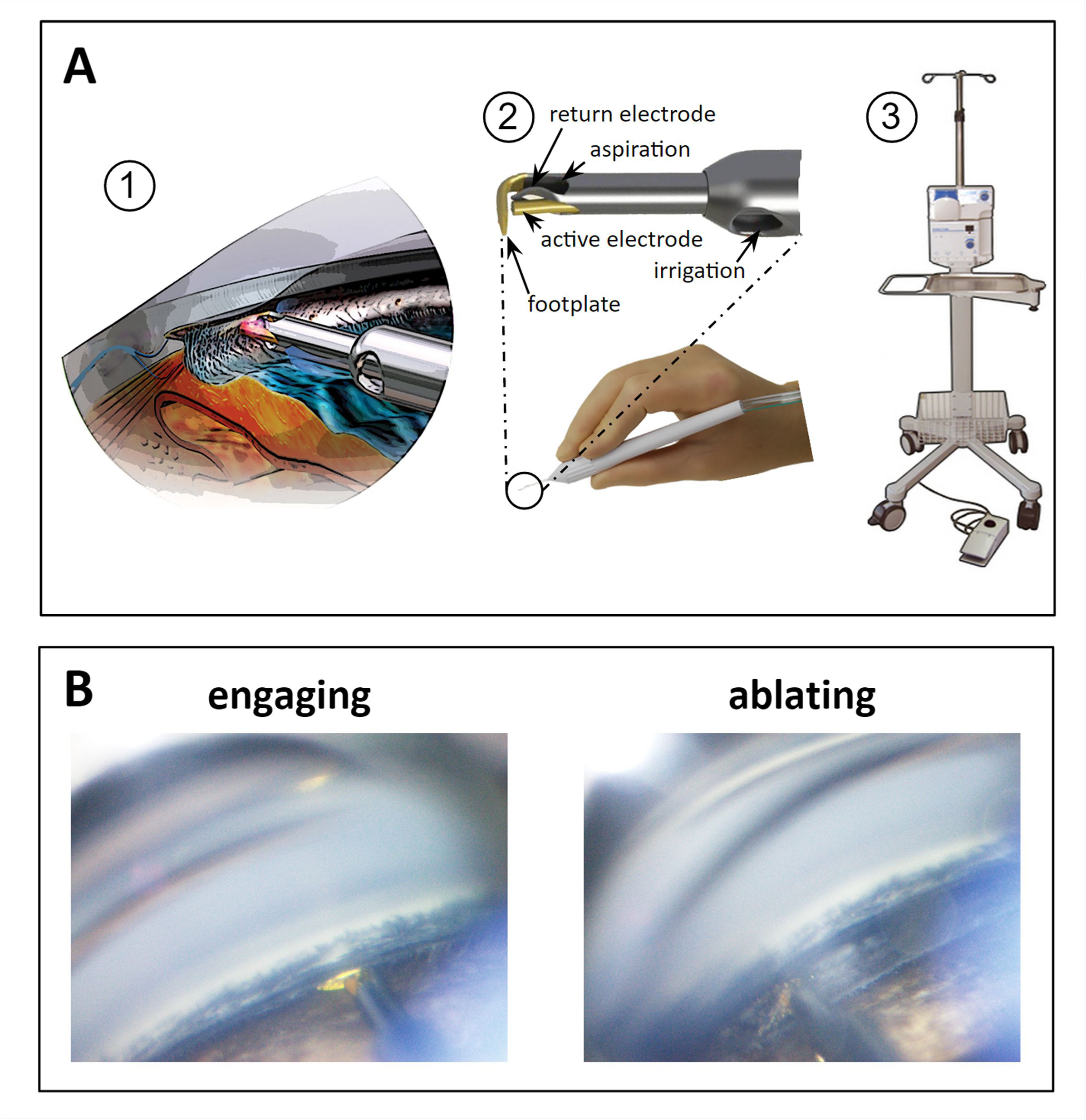
Trabectome-mediated ab interno trabeculectomy in porcine eyes. A) (1) Trabectome inserted through a clear corneal incision ablates TM that is engaged in between footplate and bipolar electrodes. (2) Handpiece and magnified view of tip. (3) Stand, operating console with peristaltic pump and high frequency generator and footswitch. B) Direct, gonioscopic view of ablation in porcine eye immediately before engaging the TM (left) and with tip obscured by TM during ablation (right).

### Histology

Segments were removed from the perfusion dish, rinsed in PBS, cut into quarters and fixed with 4% paraformaldehyde and PBS for 48 hours before being placed in 70% ethanol. A corneoscleral wedge was taken from the lateral side near the incision and one from the nasal side where the ab interno trabeculectomy had been performed (Figure 2). The section was paraffin-embedded for histological processing, cut at 6 μm thickness, and stained with hematoxylin and eosin (H&E).

**Figure 2:**
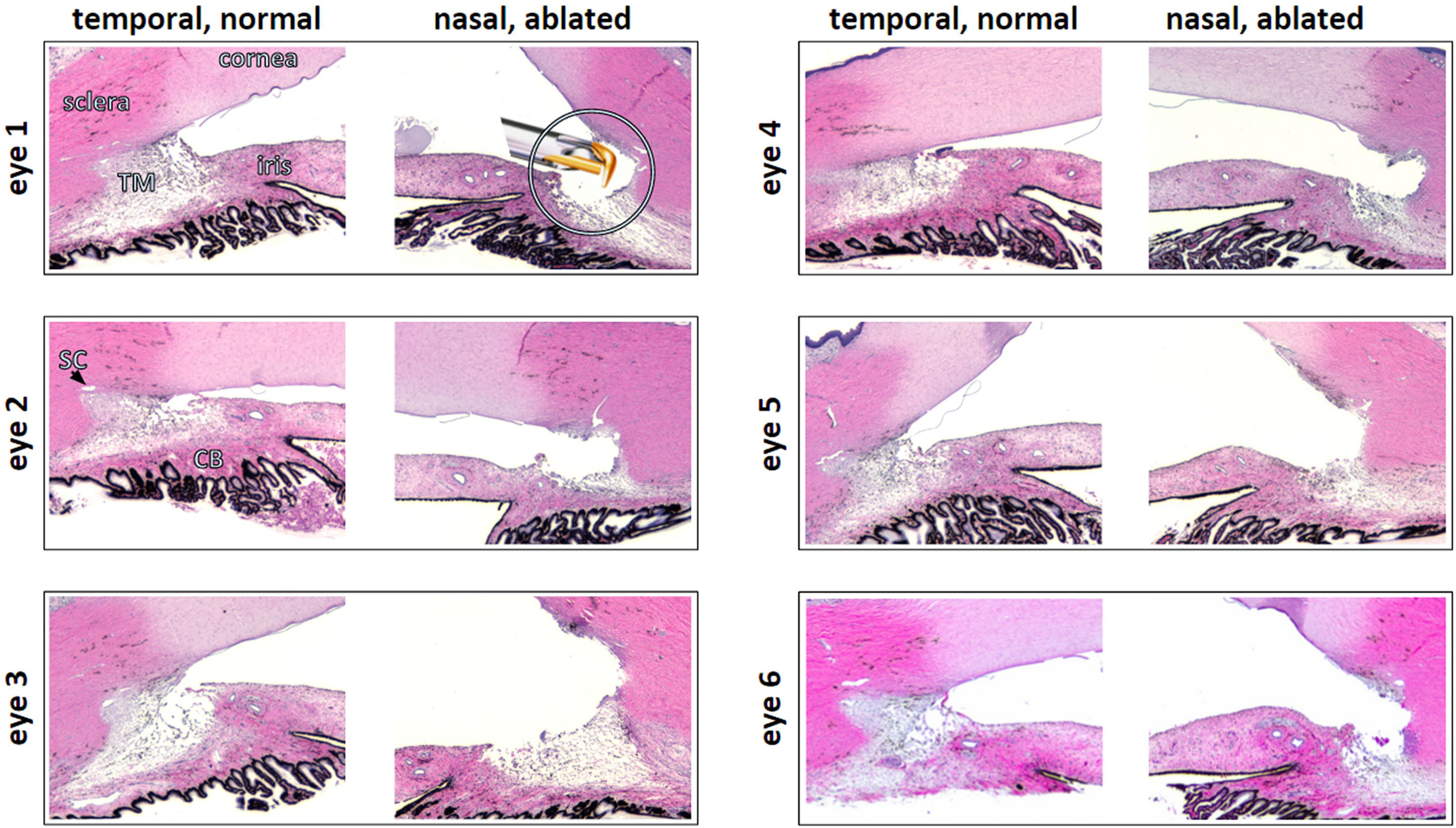
Histology of ablated, nasal TM and normal, temporal TM. Eye numbers match eyes in other figures throughout manuscript. Insert: trabectome tip shown to scale. TM= trabecular meshwork, SC= Schlemm’s canal, CB= ciliary body.

### Microsphere Canalograms

Fluorescent microsphere canalograms were obtained in six eyes. Microspheres (FluoSpheres Carboxylate-Modified Microspheres, 0.5 μm, yellow-green fluorescent (505/515), 2% solids, Thermo Fisher Scientific, Eugene, OR) were tested for a size that would not pass through the TM (Figure 3, A) while allowing visualization of the collector channels (Figure 3, B). The fluorescent microspheres were diluted 100-fold with phenol red free Dulbecco’s modified Eagle’s media (DMEM) to make the perfusate. Following the intervention described below, ab interno trabeculectomy (AIT), a 30 gauge needle was inserted through the nasal cornea 2 mm anterior to the limbus with the tip of the needle positioned in the center of the anterior chamber. Flow was began and gravity-based infusion ensued. Fluorescence was visualized with a stereo dissecting microscope equipped for fluorescent imaging (Olympus SZX16 with GFP filter cube and DP80 Monochrome/Color Camera; Olympus Corp., Center Valley, PA). Images were acquired every 20 seconds for 15 minutes for time lapse analysis (CellSens, Olympus Life Science, Tokyo, Japan) with a resolution of the 2x2 binned image capture of 580 x 610.

**Figure 3:**
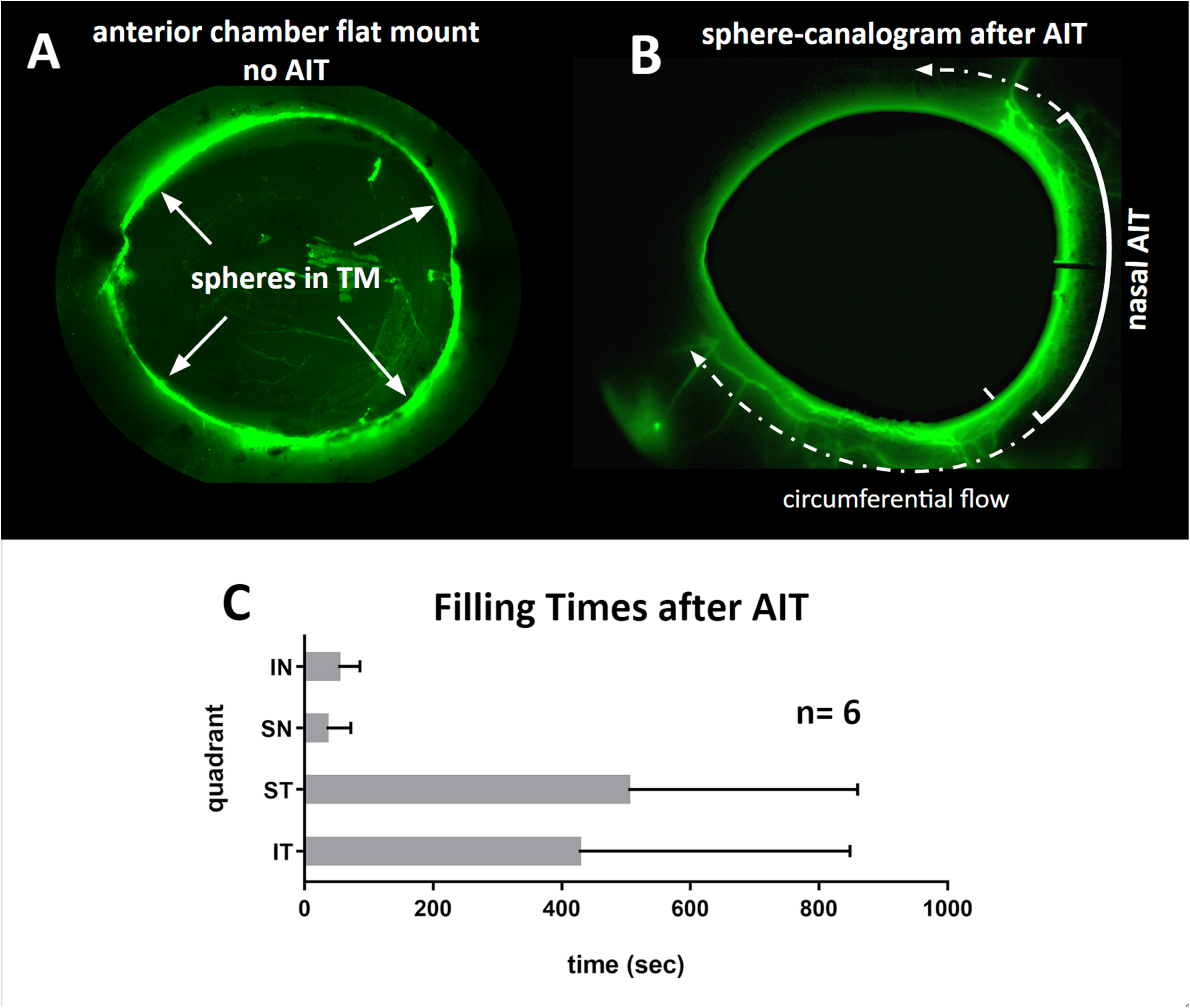
Microsphere canalograms before and after ab interno trabeculectomy (AIT). A) Perfusion of eye with fluorescent microspheres that cannot pass through the TM in a control eye (top left, inside view after removal of uvea and iris) but light up the nasal collector channel and beyond due to circumferential flow. B) Preferentially nasal outflow system filling is observed (anterior chamber subtracted using baseline). Circumferential filling can be seen (see also movie in Supplemental File 1). C) Short filling times after AIT nasal quadrants with occasional flow into adjacent quadrants. ST and IT filling times without non-filling eyes was 67±22 seconds and 46±29 seconds, respectively.

### Differential Canalograms

A fluorescent tracer reperfusion technique was used in six eyes to compare outflow changes before and after AIT detailed below using a quantitative canalography method we recently developed.^17^ Whole pig eyes were prepared and mounted under a surgical microscope as previously described. A 30 gauge needle was inserted through the nasal cornea as previously described and used to remove 0.2 mL of anterior aqueous humor from the whole eye’s anterior segment. A 30 gauge needle was then placed in the anterior segment using the exact same entry site and media with fluorescein (AK-FLUOR 10%, Fluorescein injection, USP, 100 mg/ml, NDC 17478-253-10, Akorn, Lake Forest, IL) at a concentration of 0.017 mg/ml was infused via gravity. The outflow pattern was imaged every 20 seconds for 15 minutes using a stereo dissecting microscope equipped for fluorescent imaging (Figure 4). Fluorescein flow was then stopped and the needle was removed. Following AIT, a new 30 gauge needle was placed in the anterior segment using the same entrance wound, and media with 0.28 mg/mL Texas Red (Sulforhodamine 101 acid chloride, 10 mg, Alfa Aesar, Ward Hill, MA) was subsequently infused via gravity. Again the outflow pattern was imaged every 20 seconds for 15 minutes. The eyes were then fixed and sent for histology.

**Figure 4:**
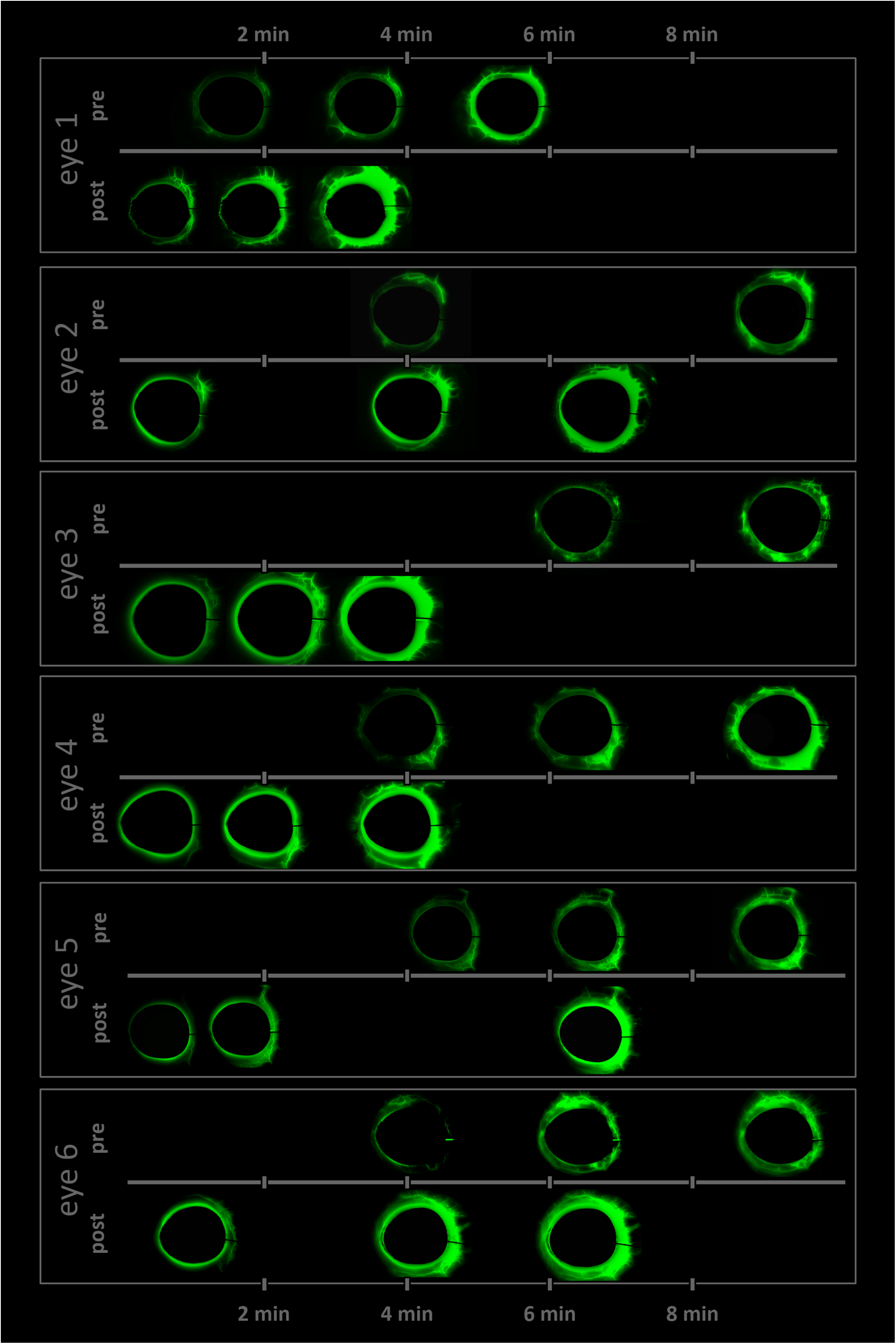
Time lapse of differential canalograms for each eye. Images are subtracted from baseline to suppress nonspecific fluorescence. Frames selected show filling of new segments. Pre-and post-ab interno trabeculectomy canalogram frames are matched (also see movie for supplemental File movie 1).

To determine differences in chromophore detection sensitivity, a hemocytometer chamber was filled with 10 μL each of Fluorescein and Texas Red at the previously stated concentrations. Proper exposure times of 15 ms for Fluorescein and 10 ms for Texas Red were determined with the fluorescence equipped stereo dissecting microscope above.

Initially, six eyes were perfused with fluorescein first immediately followed by Texas Red without AIT. Another six eyes were perfused with Texas Red first followed by fluorescein to determine whether the order of perfusion would have any effect. When there was no statistically significant difference in filling times between the two (p=0.06, n=12), but a 19% slower filling rate for Texas Red, the order was chosen to be fluorescein followed by Texas Red for all eyes in order to avoid false positive flow enhancement by AIT.

### Quantification of Outflow Change

As described before,^17^ we used a program written in R^18^ to automatically compute the focal outflow changes (Figure 5, left) and convergent perilimbal aqueous flow (Figure 5, right) using the eye-canalogram package to process the image datasets, the source code of which we made available for download (R package “eye-canalogram”, https://github.com/enbrown/eye-canalogram/tree/06461498c8).

**Figure 5:**
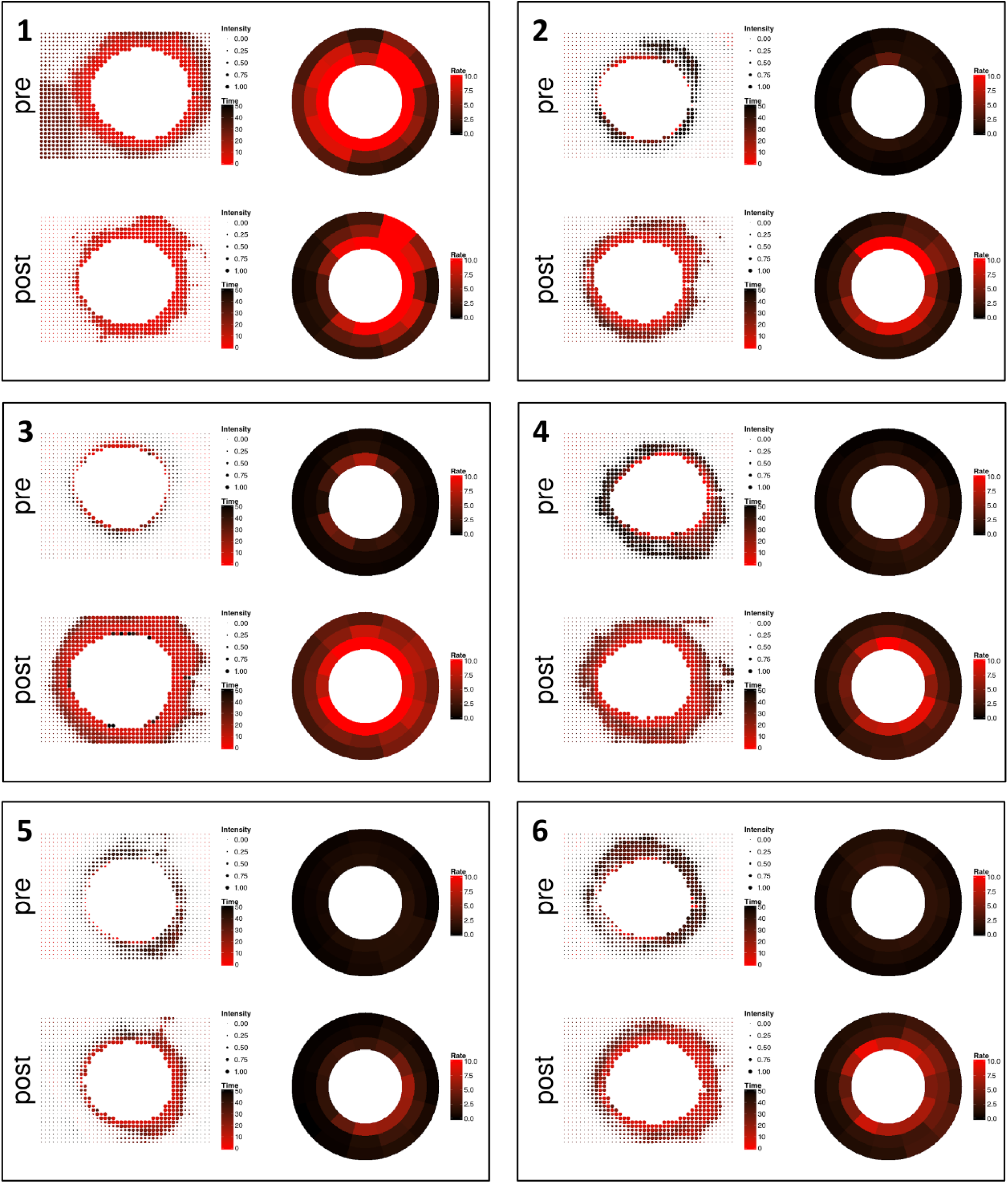
Quantitative canalograms with individual intensity fits (left) and circumferential flow rates (right). Both are increased in most eyes, even in quadrants away from the nasal AIT site.

Briefly, for each image set, the cornea was first manually segmented. The image resolution was then reduced to 32 x 32 macropixels per image. Generalized additive models (GAM) were fit for each macropixel. Metrics from the fit were recorded and graphically displayed (Supplemental File 2). In dot plot images, larger pixels corresponded to more intense fluorescein signals while red dots corresponded to faster filling. In addition, all regions (clock hour and three radial rings) were fit to a single GAM using smoothing terms for the radial ring, clock hour, and frame number. Pre-and post-treatment image sets for each eye were registered using the clock hour and radial distance to compute the change in fit metrics (Figure 6A). Similarly, all eyes were warped to a common reference frame and averaged to produce Figure 6B.

**Figure 6:**
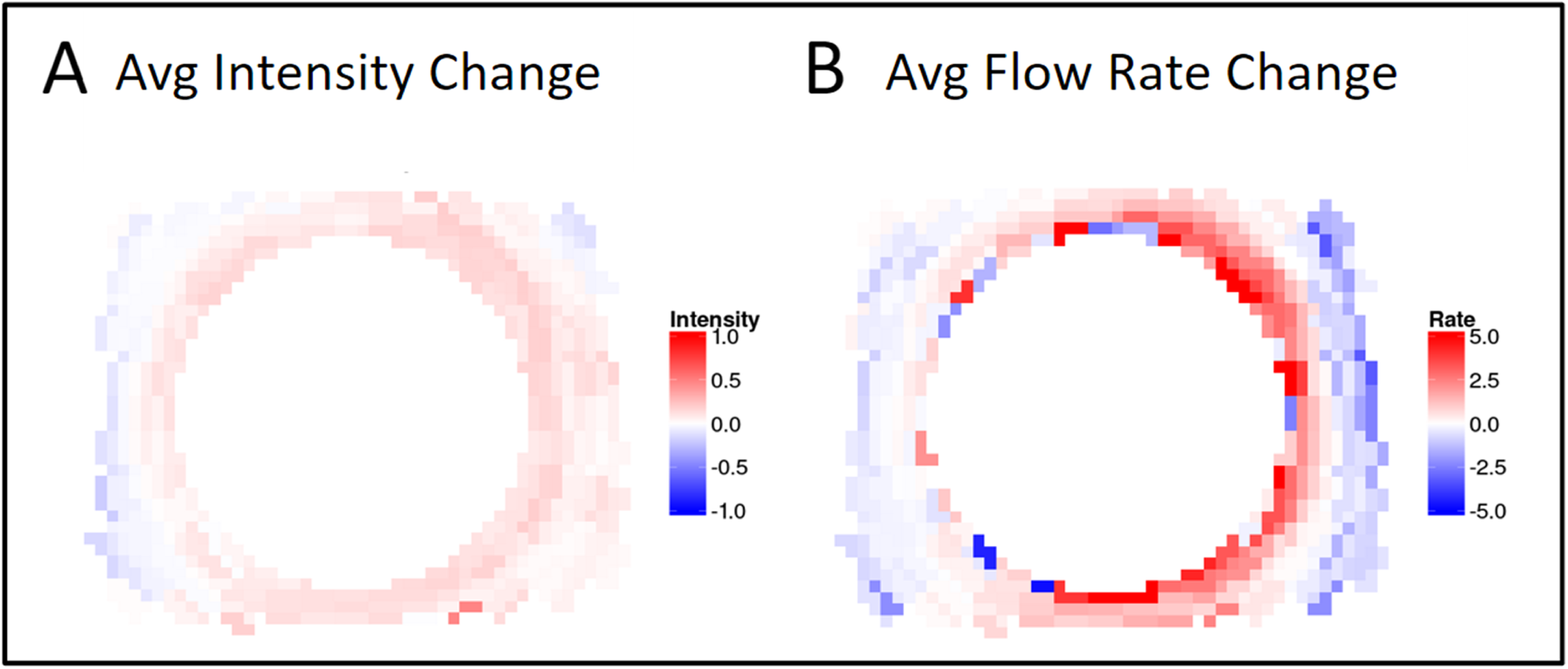
A) Quantitative change analysis AIT. Each eye produces a pair of images: change in fluorescence intensity (left image) and change in rate of fluorescence uptake (right image). Red (positive values) indicates greater intensity or rate of uptake following AIT; blue (negative values) indicates less intensity or reduced uptake. B) Summarizing graph of intensity and rate change for all six eyes.

Each quadrant was also analyzed for first detection of fluorescence in an outflow structure downstream of Schlemm’s canal segments. Timestamps from time lapse recordings were summarized as graphs (average ± standard deviation (SD)) for the inferonasal (IN), superonasal (SN), superotemporal (ST), and inferotemporal (IT) quadrants.

## Results

Visualization and TM ablation could be achieved with a standard trabectome system and the kit-included, modified Swan Jacob goniolens (Figure 1, A). Pectinate ligaments had to be gently lysed (Figure 1, B, left) before the TM could be engaged for ablation (Figure 1, B, right). Power and aspiration settings were identical to surgery in human patients. During ablation, the footplate encountered more stops than typical for human eyes when SC segment ends were reached.

The histological analysis showed that the relatively prominent TM and common SC segments had been ablated with a few exceptions of SC segments too small to enter with the trabectome tip (Figure 2, eye 1, shown to scale). No coagulative damage was observed.

Eyes without AIT that were infused with fluorescent spheres showed fluorescence only in the TM. When those were hemisected and vitreous and iris removed, sphere distribution appeared relatively even throughout the TM circumference (Figure 3 A). In contrast, canalograms with fluorescent spheres obtained after AIT experienced fast filling of proximal and distal parts of the outflow system along the ablation site followed by circumferential and then centrifugal filling of adjacent quadrants. Filling of nasal quadrants was nearly ten times faster nasally than that of temporal quadrants (Figure 3 B).

Differential canalograms, obtained before and after AIT (Figure 4 and Supplemental File 1: Movie with parallel canalograms pre-and post-AIT) had a 17±5 fold increase in filling inferonasally, 14±3 fold increase superonasally and also an increase in the adjacent quadrants with a 2±1 fold increase superotemporally and 3±3 inferotemporally. The superonasal quadrant was the fastest to fill (p<0.5) followed by the inferonasal and super-and inferotemporal quadrants (Figure 4 and Figure 7). Although fluorescent dyes used in these differential canalograms were not blocked by the TM like the fluorescent spheres, superotemporal and inferotemporal quadrants often filled circumferentially from the site of AIT.

**Figure 7:**
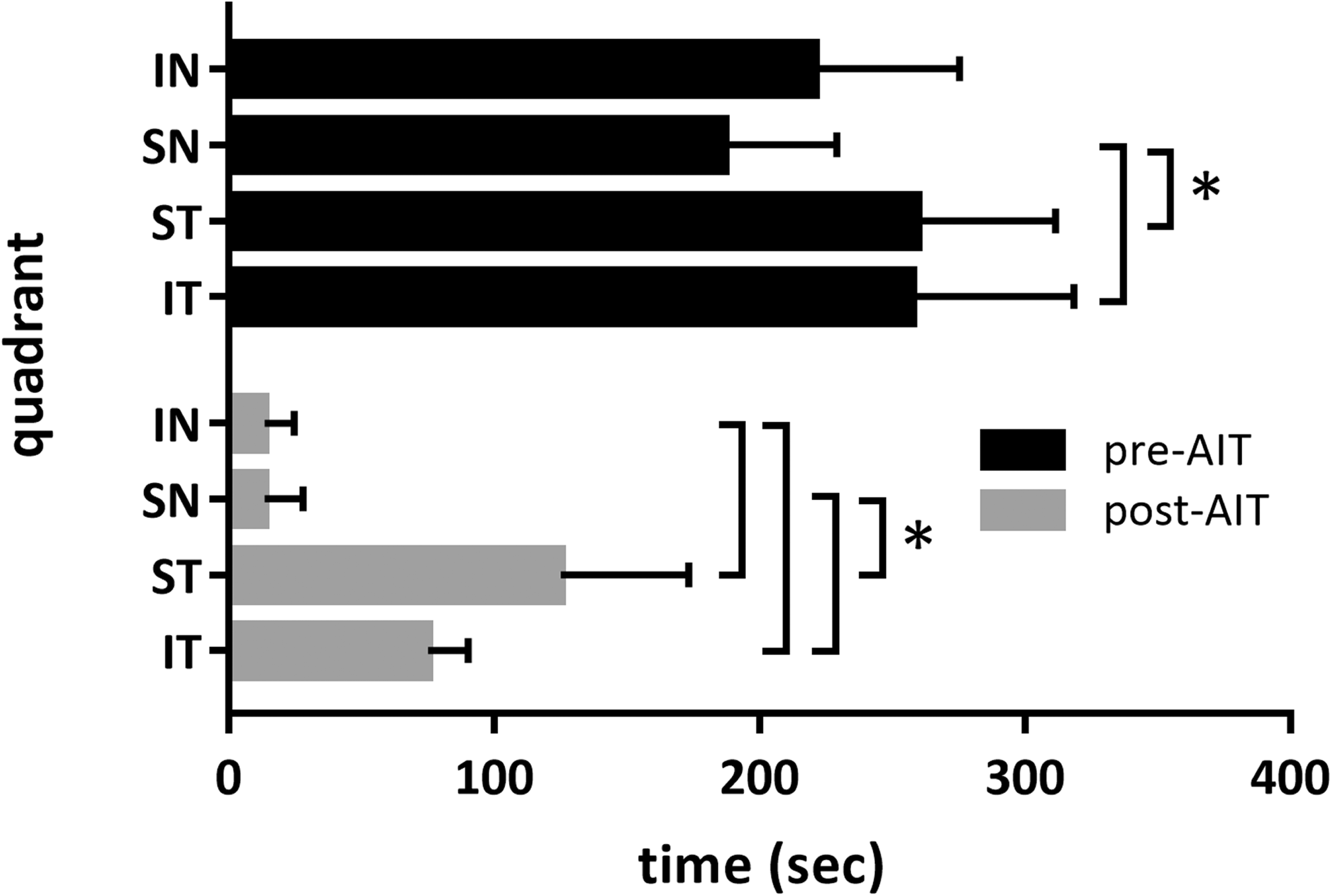
Outflow change summary after AIT. Short bars indicate fast filling not only in the nasal quadrants (IN: inferonasal, SN: superonasal) but also in the temporal quadrants (ST: superotemporal, IT: inferotemporal) following AIT.

Individual fit curve analysis indicated a shift in filling speed towards the nasal side and an overall faster filling throughout the remainder of the outflow system in most eyes (Figure 5). This can be seen by a shift towards red in the color coded analysis indicating a shorter time towards peak fluorescence, as well as the increased bubble size that describes the peak fluorescence obtained. When perilimbal flow was examined specifically, circumferential filling was evident. With the exception of eye 1, all eyes post-AIT showed an initial uptake centered on the inferonasal and superonasal quadrants as expressed by the red color coding that spreads circumferentially from here (Figure 5).

Changes from pre-to post-AIT fluorescence intensity changes are summarized in Figure 6. In average of all six eyes, peak intensity increased in the inferonasal quadrant and in the superonasal quadrant as well as in the remainder of the circumference (Figure 6 left). In contrast, flow rate changes from pre-to post-AIT are more substantial in the nasal quadrants.

These changes from pre-to post-AIT were statistically significant in all quadrants but more pronounced in the inferonasal and superonasal quadrant. Those quadrants remained significantly faster than the superotemporal and inferotemporal quadrants. There was no difference between the superotemporal and inferotemporal quadrant before or after AIT.

## Discussion

Focal changes of conventional aqueous humor outflow are difficult to examine^8,19^ yet are crucial to better understand how elements downstream of the TM may influence outflow. The lack of an inexpensive, readily available and high quality outflow model has impeded glaucoma research and training of surgery on this submillimeter structure alike. Here, we address both with a differential canalography technique that allows to examine the effect of trabectome-mediated AIT on focal outflow in porcine eyes.

Visualization of the chamber angle and ablation of TM by AIT was quite similar to surgery in human patients. Angle surgery can be practiced in human eyes^20^ but costs of those are high and the corneal clarity is often too compromised to see the angle.^21^ As a result, artificial eyes were preferred by trainees.^21^ The histology following AIT shows that despite the anatomical differences between the pig and human chamber angle, an extensive ablation of TM can be achieved without an obvious coagulative thermal effect on adjacent tissues. Different from coagulation devices, the instrument used here generates plasma to ionize tissue and has a highly confined heat dissipation cone that is similar to photodisruptive lasers.^11^

The initial canalograms obtained with fluorescent microspheres demonstrated that TM must be removed before they can enter the conventional outflow system. Although the angular aqueous plexus of pigs does not have a continuous Schlemm’s canal as primate eyes do, but rather multiple SC-like segments, we observed circumferential flow that extended far beyond the ablation site. This indicates that canal segments are connected and that supraphysiological flow from the site of ablated TM is displacing the normal flow that is still occurring through the non-ablated TM.

The perilimbal flow analysis of differential canalograms showed high flow areas near the inferonasal and superonasal angle. This matches the areas in between the recti muscles, where larger collector channels reside. The circumferential flow patterns observed here contradicts the assumption of noncontinuous SC segments in the pig. A three dimensional reconstruction at a higher resolution than conventional histological sections by SD-OCT may be necessary. Such an approach has enabled discovery of previously overlooked valve-like elements in human eyes.^22,23^

We provide a heatmap summary image that combines fluorescence and flow rate of all six eyes. This allows to visualize how flow can be enhanced in non-glaucomatous eyes to levels above the physiological flow rate and beyond the TM ablation area. In this constant pressure perfusion system AIT led to increased flow and peak fluorescence in all quadrants, beyond the nasal site where AIT was performed. This has practical implications for patient care and suggests that it may not be necessary to obtain a very extensive ablation or circumferential access to the outflow tract. Carefully observing glaucoma surgeons have previously described fluid waves of saline displacing blood in collector channels after AIT.^24^ These appeared to be limited to the site of ablation when observed through an operating microscope. Such visualization may relatively underestimates the amount of flow that could be detected with more sensitive, fluorescent dyes as used here.

There are well established angiography methods for organs larger than the anterior segment of the eye, such as the heart.^25^ Characteristic for such angiography is that vessels branch off large primary vessels to subsequently smaller ones. The outflow tract of the eyes is different and has a much more diffuse and connected nexus of vessels with variable caliber. The drainage system just distal to the outer wall of the SC is more similar to the honeycomb pattern of a capillary network but its vessels are larger and collapsible depending on the perfusion pressure. This makes it challenging to use Doppler strategies to compute flow speeds in single vessels. The data presented here validates the use of fluorescence change^17^ as a surrogate to tracking reflective particles.

In conclusion, we developed a method to quantify outflow enhancement from plasma-mediated ab interno trabeculectomy and show that nasal ablation of trabecular meshwork increased outflow not only locally but circumferentially. Methods and code provided here will aid further investigations into segmental outflow changes.

